# Chemotactile perception and associative learning of amino acids in yellowjacket workers

**DOI:** 10.1101/2023.12.16.572014

**Authors:** Analía Mattiacci, Ana Laura Pietrantuono, Maité Masciocchi, Juan C. Corley

**Affiliations:** Grupo de Ecología de Poblaciones de Insectos, IFAB-CONICET-INTA EEA Bariloche. Modesta Victoria 4450. San Carlos de Bariloche (8400), Río Negro-Argentina; Departamento de Ecología, Centro Regional Universitario Bariloche, Universidad Nacional Del Comahue, Bariloche, Argentina

**Keywords:** differential conditioning, MaLER, social wasps, Vespula

## Abstract

Learning and memory are essential for animal survival, influencing preferences, decision-making, and foraging behaviour. In this study, we explore the perceptual and learning abilities of *Vespula germanica* (yellowjacket wasps) to various amino acids. We hypothesize that *V. germanica* can qualitatively evaluate various amino acid solutions, given their scavenging habits and the possibility of metabolizing amino acids to fuel energy. Through chemo-tactile differential conditioning, we studied worker wasp’ maxilla labium extension response (MaLER) to essential (Lysine, Tryptophan, Arginine) and non-essential amino acids (Ornithine, Aspartic acid, Glycine). Conditioning sessions included individual amino acids against water and comparisons between different amino acids. Additionally, we tested retention, discrimination, and generalization abilities, 30 minutes later with conditioned and novel stimuli. Our results show that wasps exhibit the ability to learn and discriminate various amino acids. The discrimination capacity extended to differentiating between pairs of amino acids. Memory retention was generally robust, but certain associations observed during conditioning did not persist after a 30-minute interval. Moreover, when wasps were trained with essential amino acids, the acquired learning did not generally extend to other non-essential amino acids, except for Arginine, which exhibited generalization when tested with its precursor, Ornithine. Conversely, when trained with non-essential amino acids, the acquired learning generalized to other essential amino acids. These results suggest that, unlike other hymenopterans, wasps can detect, discriminate, and generalize free amino acids, crucial for their foraging decisions. This knowledge contributes to understanding the cognitive dimensions of *V. germanica* and their implications for targeted pest management.

**Summary statement:** This study on *Vespula germanica’*s amino acid perception enhances our understanding of foraging behaviours. The findings contribute to insect cognition studies, with potential implications for pest management.

## Introduction

Learning -the acquisition of neuronal representations of new information- and memory -the retention of newly acquired information over time-are critical processes for many animal species since they influence their preferences, decision-making, and foraging behaviour, ultimately determining fitness success (Dukas, 2008). During foraging, animals must actively search for resources such as food to obtain essential dietary macro-nutrients (i.e., carbohydrates, protein, and lipids; Bell, 1990). Evaluating the nutritional quality of food is complex due to the diverse array of nutrients and compounds present. Olfactory cues, like volatile organic compounds, aid in locating food from a distance, while taste, or contact chemoreception, is crucial for non-volatile nutrients (such as proteins: Chapman, 1998). Yet, compared to olfaction, there is significantly less understanding of chemotactic sensation, especially in invertebrates. Understanding the sensory, perceptual, and cognitive capacities of organisms involved in this process is important for understanding foraging decisions.

Protein consumption (primarily essential amino acids which are those that animals cannot synthesize) is crucial for various functions, including protein synthesis, enzymatic activity, transportation, and storage, as well as their role as neurotransmitters. Animals need to break down ingested proteins to obtain or synthesize amino acids. Thus, internal monitoring of amino acid demand and the organization of behaviour to secure their supply is important to any heterotopous organism. Despite their importance for individual development and reproduction, how amino acids are perceived and how organisms forage for them remains an open question (Ruedenauer et al., 2019).

In social insects (such as ants, bees, and wasps) the metabolic requirements of all individuals in the colony depend on food searching performed by foraging individuals (Perry, 1984). All colony members require the same set of 20 amino acids crucial for constructing proteins in all living organisms (Klowden, 2013). In most social insects, the proteins collected are mainly for feeding larvae, except in some species of vespids. It has been noted that the presence of intestinal proteases in *V. germanica* and *Vespa orientalis* workers allows digestion and metabolization of proteins obtained in the absence of larvae (Grogan and Hunt, 197; Bodner et al., 2022). Indeed, recent studies showed that *V. vulgari*s and *V. orientalis* workers could use some amino acids as metabolic fuel (Teulier et al., 2016). Therefore, the success of the colony relies heavily on efficient foraging strategies performed by workers and their ability to perceive, learn, and remember relevant environmental cues.

From a cognitive perspective, one of the most studied social insects is the honeybee *Apis mellifera* (Bitterman et al., 1983; Giurfa and Sandoz, 2012). These studies on honeybee learning have been performed by observing the proboscis extension response (PER). This innate response occurs when honeybees land on flowers and detect the presence of nectar with their antennae. Thus, individuals can associate flower shape, colour, and odours with the presence of food; in other words, associative learning is induced. Similarly, wasps and ants extend their maxilla-labium apparatus (i.e., maxilla-labium extension response or MaLER) allowing us to study associative learning (Guerrieri and D′Etorre, 2010; Mattiacci et al., 2021). In classical associative learning protocols, the goal is to establish an association between an unconditioned stimulus (US) like the sucrose solution, and a conditioned or neutral stimulus (CS) like an olfactory or chemotactic cue. If individuals successfully learn this association, they will, upon repeated exposure, extend their proboscis or maxilla-labium complex in response to the mere presentation of the CS (Bitterman et al., 1983; Matsumoto et al., 2012). According to the properties and characteristics of the resources, insects also can learn to discriminate or generalize stimuli. Discrimination allows individuals to distinguish different stimuli, whilst generalization allows them to classify similar, though different stimuli, into the same perceptual category (Getz and Smith, 1987; Laska et al., 1999; Gumbert, 2000; Ghirlanda and Enquist, 2003; Guerrieri et al., 2005; Robazzi Bignelli Valente Aguiar et al., 2018; Pietrantuono et al., 2019).

*Vespula germanica* (Hymenoptera: Vespidae; Fabricius, 1793), commonly known as “yellowjacket wasps”, is a social species that has successfully invaded and become a pest in various regions worldwide, including southern Argentina. During adulthood, yellowjackets are a generalist and opportunistic species that forage on diverse food resources. These wasps obtain carbohydrates from any available source of sugars, whether natural or of anthropogenic origin, while carrion is the main source of lipids and proteins (Raveret Richter, 2000; Pereira et al., 2016). Previous studies showed that free-flying forager wasps of *V. germanica* exhibit strong learning abilities: they can associate specific visual, spatial, and olfactory cues with resource-rich food sources (D’Adamo & Lozada, 2008; Lozada and D’Adamo, 2011). However, to date, limited studies address the perceptual and cognitive capabilities of *V. germanica* under laboratory conditions (but see Mattiacci et al., 2021, 2023; Masciocchi et al., 2023).

We aimed to investigate the perceptual and learning abilities of *V*. *germanica* to various amino acids. Specifically, we seek to understand whether these wasps can perceive, learn, discriminate, and generalize information about amino acids based on their sensory experiences, focusing on antennae-driven response under a MaLER chemotactic differential conditioning protocol. Given the relevance of specific amino acids for nutrition and the capacity of wasps to digest and utilize amino acids as an energy source, we hypothesize that *V. germanica* can evaluate qualitatively a wide array of amino acid solutions. Specifically, we determined whether wasps are able to (1) perceive some specific amino acids through chemoreception in their antennae, (2) perform associative learning toward amino acids, (3) discriminate among stimuli of different molecular and nutritional profiles, and (4) generalize among stimuli. The results suggest that *V. germanica* wasps can detect free amino acids. In a natural foraging context, this may translate into an ability to differentiate and learn between two food sources with different nutritional qualities. The information obtained in this study provides information that may be useful for the management (currently mediated mainly by toxic food-based baits) of this highly invasive species.

## Materials and methods

### Wasp capture and harnessing

We captured foragers of *V. germanica* during March and May of 2021 and 2022 (peak flight period) from different sites in the proximities of the city of San Carlos de Bariloche, Río Negro, Argentina. We randomly offered feeders with 15 g of ground beef at eight different places (between 9 AM and 1 PM). We caught wasps as they landed on the feeder and kept them in individual plastic containers (5 cm height x 8 cm diameter), we captured around 30 individuals daily. We immediately transferred all captured wasps to the laboratory to carry out the experiments. We kept the wasps under controlled conditions with low temperatures (5° C) and darkness to induce a decrease in the metabolism until the beginning of experiments (Käfer et al., 2012). After this resting period, we anesthetized individuals using a brief exposure of approximately 10 seconds to carbon dioxide. This minimal exposure duration facilitated swift experimental manipulation and rapid recovery. During this lapse of time, we placed each individual into a modified Eppendorf tube (5 ml), with the bottom cut off to allow the wasp’s head to be gently pushed through the hole. We harnessed each wasp with tape on the back of the head, allowing free movements of the antennae and mouthparts (Fig. 1A; see Bitterman et al., 1983). After 1 hour of recovery from the anaesthesia and before beginning the learning protocol, we tested the motivation of each individual. We determined that an insect was suitable for the experimental procedure when it responded to a sucrose solution 50% v/v -by touching each wasp’s antennae with a toothpick- and producing the maxilla labium extension response (MaLER, Fig. 1B; Pietrantuono et al., 2019; Mattiacci et al., 2021). Individuals that did not responded were discarded from experiments. To avoid any effect of external odours, we placed the Eppendorf rack with the harnessed wasps inside an air extraction hood.

**Figure 1.**
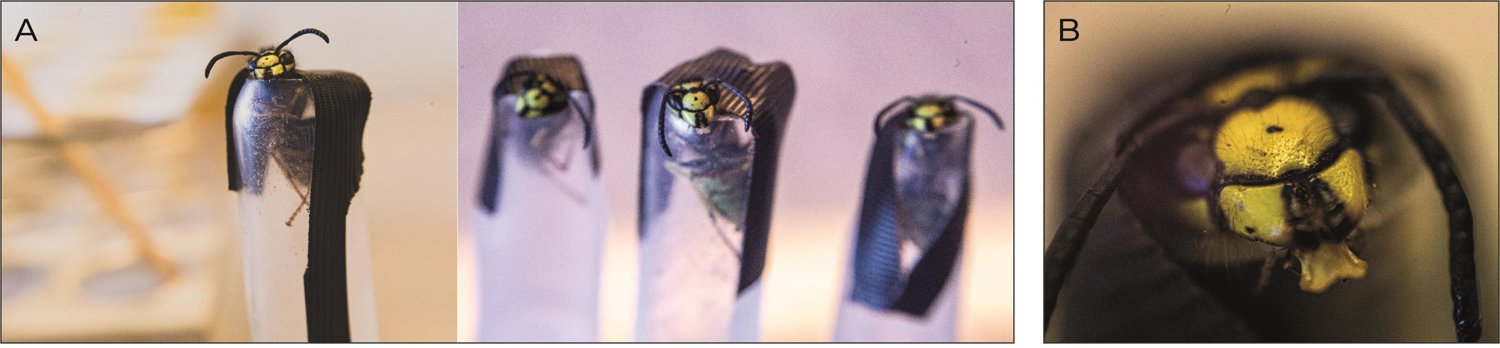
Experimental Set-Up for Individual Perception and Learning in *Vespula germanica* Wasps using MaLER Technique. Wasps harnessing A) ready to start the learning protocol and B) performing the maxilla labium extension response (MaLER). The experimental setup utilized the MaLER technique to explore individual perception and learning in a chemotactic differential conditioning protocol. Each harnessed wasp, confined to a plastic tube, retained free movement of antennae and mouthparts. The protocol assessed perceptual and learning abilities by stimulating restrained individuals’ antennae with a toothpick carrying one solution paired with a reward, while another solution remained unrewarded.

### Preparation of stimuli

We used as gustative stimuli six amino acids (three essential and three non-essential amino acids) with different molecular profiles and physiological roles. The three essential amino acids were: (1) Lysine (Lys), crucial for memory consolidation, and found in larval saliva; (2) Tryptophan (Trp), present in wasp larval saliva; (3) Arginine (Arg), pivotal in memory consolidation in bees, serving as a precursor to ornithine. While the three non-essential amino acids were: (4) Ornithine (Orn), contributing to memory consolidation in bees; (5) Aspartic acid (Asp), playing a fundamental role in synthesizing Arginine and Lysine; (6) Glycine (Gly), a metabolic fuel in hornet, is present in wasps larval saliva (Hunt et al., 1982; Lopatina et al., 2017; Pratavieira et al., 2019; Marchi et al., 2021; Bodner et al., 2022). For the learning protocol (see below), we dissolved all amino acids in deionized water at a concentration of 10 mg/ml.

### Learning protocol

To determine whether wasps can perceive, learn, and discriminate or generalize the amino acids, we used a chemo-tactile differential conditioning protocol between two conditioned stimuli, one rewarded (from now on CS+) with sucrose solution (unconditioned stimuli or US) and the second one, unrewarded (from now on CS−). Employing a differential conditioning approach, where only one of two different stimuli is rewarded, allows us to evaluate whether individuals can differentiate between these two stimuli. We carried out two different conditioning procedures with two different aims:

#### 1. Conditioning A: Amino acid vs water

We evaluated each amino acid against pure water to see if wasps could perceive it. It is worth noting that pure water is a neutral stimulus for non-thirsty wasps since they do not exhibit MaLER (unlike honeybees; Page et al., 1998; Mattiacci et al., 2023). We conditioned a total of 312 forager wasps distributed on 12 different conditioning sessions for the amino acid as a positive or negative stimulus (*e.g.,* Glycine as CS+ and water as CS-, or vice versa).

#### 2. Conditioning B: Amino acid vs amino acid

We then evaluated amino acids against each other to see if the wasps could differentiate them. Specifically, we evaluated one essential amino acid against a non essential one, for two pairs of amino acids (e.g., Lysine as CS+ and Aspartic Acid as CS-, Tryptophan as CS+ and Glycine as CS-; or vice versa) We conditioned a total of 97 forager wasps distributed in 4 conditioning sessions.

The learning protocol consisted of two phases: the *Conditioning phase* and the *Test phase* (protocol adapted from Ruedenauer et al., 2019; Pietrantuono et al., 2019). It should be noted that we assume a positive response to the offered stimulus when an individual performs a complete MaLER reflex.

### Conditioning phase

For the *Conditioning phase*, we placed one individual in a rack (one wasp per experimental series) and allowed it to rest for 20s. Immediately after, we presented the stimulus and touched an antenna for 10s with a toothpick soaked in the solution (CS). The wasp could freely sense the stimulus with the tip of its antenna. Five seconds after stimulus onset, we presented it with another toothpick touching the free antenna, either soaked with sucrose solution (as US) or clean (to equalize visual cues for the CS+ and CS− presentation). If the wasp extended its maxilla labium complex, we allowed it to lick on the toothpick. We removed the US and CS together after 5s. Subsequently, we allowed the wasp to rest for another 5s before being replaced by the next. The time between trials (inter-trial interval, ITI) was between 5 and 10 min. The number of trials was 12 per individual (six for CS+ and six for CS−) presented in a pseudo-randomized order (Fig. 2). We used 10 different pseudo-randomized sets of trials randomly assigned to each individual (for example, CS-, CS-, CS+, CS-, CS-, CS+, CS+, CS-, CS-, CS+, CS+, CS+). We tested each stimulus once as CS+ and once as CS− (reversed meaning), and each animal only once.

**Figure 2.**
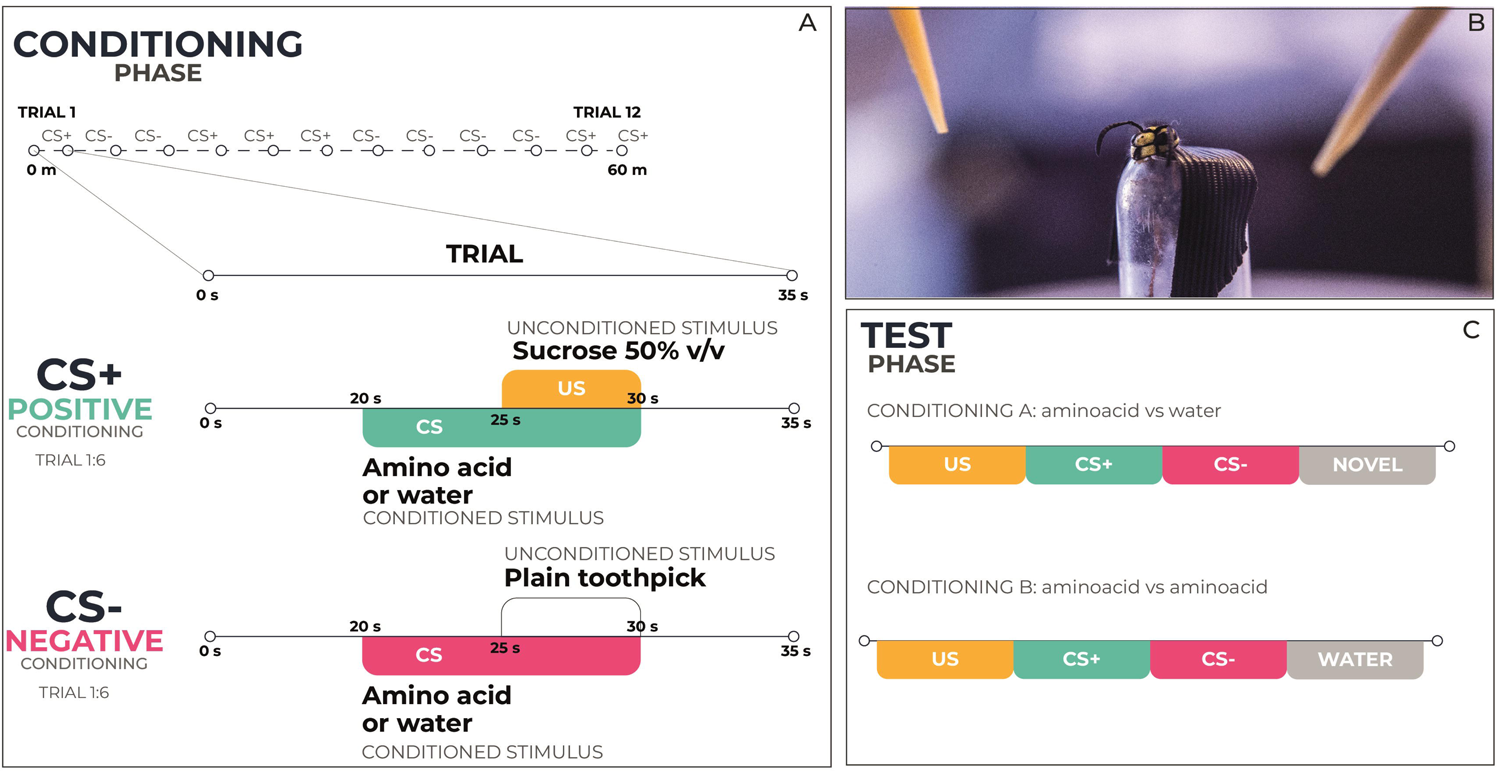
Learning Protocol in *Vespula germanica* Foragers - Conditioning and Test Phases. (A) During the *Conditioning phase*, we subjected each wasp to 12 trials (six for CS+ and six for CS−) in a pseudorandomized order. Each trial lasted 35 seconds. First, the wasp rested for 20 seconds in a rack. We then introduced a stimulus by touching one antenna for 10 seconds with a toothpick soaked in a solution (CS) rich in amino acids or pure water, according to the conditioning session. The wasp could freely sense this stimulus with the antenna’s tip. Five seconds later (i.e., 25 seconds from the beginning of the trial) we presented another toothpick to the other antenna (B). For positive conditioning (CS+, green) we presented the toothpick soaked with a sucrose solution (US). However, for negative conditioning (CS-, pink) we presented a clean toothpick (to equalize visual cues for CS+ and CS− presentation). We removed both the US and CS after 5 seconds (i.e., 30 seconds from the beginning of the trial). Afterward, the wasp rested for 5 seconds before being replaced by the next one. (C) During the T*est phase*, we only offered the individual four stimuli (one stimulus per trial in an unexpected way) without the reward and observed the insect response. We tested wasps trained against water (Conditioning A) with sugar (US), the conditioned amino acid (CS+ or CS-according to the conditioning session), water (CS+ or CS-according to the conditioning session), and a novel amino acid. Additionally, wasps trained against a different amino acid received tests with sugar (US), both conditioned amino acids (CS and CS-), and water (as a negative control).

### Test phase

After the *Conditioning phase* (either A or B) and an inter-trial interval of 30 minutes, the *Test phase* began. At this stage, each wasp was presented with four stimuli (one stimulus per trial) in a random sequence without the reward. When wasps were conditioned with amino acids and water (Conditioning A), the stimulus could be water, the same conditioned amino acid, a new one, or the sucrose solution (as a positive control) to assess motivation and fatigue after the procedure. Individuals who did not respond to the sucrose solution were discarded from the statistical analysis. If the wasps extend their maxilla labium apparatus in response to a novel stimulus (i.e., a different amino acid from the one used for conditioning), it indicates that the stimulus was perceived as similar to the CS+, providing evidence of generalization. In this experiment, we aimed to assess the ability of wasps to recognize the conditioned stimulus and their generalization capacity between two different amino acids. For wasps conditioned with two different amino acids (Conditioning B), we offered four stimuli during this phase (also randomly): water (as a negative control), the conditioned amino acids (to evaluate memory retention), and the sucrose solution.

### Data analysis

For the 16 conditioning sessions (12 against pure water and four among different amino acids), we analysed the probability of MaLER behaviour occurrence following a binomial distribution response variable and using a Generalized Linear Mixed Model (GLMM) with a Bernoulli error structure. We considered “conditioned stimulus” (*i.e.,* rewarded or unrewarded), and “trial” (*i.e.,* from 1 to 6) as fixed effects with their interactions. Also, we considered “individual” as a random effect. We performed comparisons between CS+ and CS-learning curve responses. Besides, for each conditioning session, we assessed MaLER behaviour occurrence for four stimuli (test phase) by GLMM with a Binomial distribution, “stimulus” as a fixed effect and “individual” as a random effect. Comparisons between each stimulus were performed. We fitted the models in the R program (R Development Core Team, 2019) using the *glmer* function of the *lme4* package (Bates *et al.,* 2014). We fitted *Post hoc* comparisons with the *glht* function of the R package *multcomp* (Hothorn et al., 2008).

## Results

### Conditioning phase

#### Conditioning A: Amino acid vs water

Wasps could learn and discriminate all evaluated amino acids against water (Lysine, Tryptophan, Arginine, Ornithine, Aspartic Acid, and Glycine; Fig. 3). Arginine and Aspartic acid triggered this behaviour when presented as CS+ (Arginine: p_Arg-Wat_<0.001, p_Wat-Arg_=0.575; Aspartic acid: p_Asp-Wat_=0.019, p_Wat-Asp_=0.736), Ornithine when presented as CS-(Ornithine: p_Orn-Wat_=0.355, p_Wat-Orn_<0.001), and Lysine, Tryptophan and Glycine in both conditions (Lysine: p_Lys-Wat_<0.001, p_Wat-Lys_<0.001; Tryptophan: p_Trp-Wat_<0.001, p_Wat-Trp_=0.075; Glycine: p_Gly-Wat_=0.008, p_Wat-Gly_<0.001). In all the curves, responses to CS+ stimuli showed an upward trend, indicating a positive response to the conditioned stimulus. We noticed differences in the learning curves between essential and non-essential amino acids. For the essential amino acids (i.e., Lysine, Tryptophan, and Arginine), whether trained as CS+ or CS-, the final trial exceeded the 50% mark (Lysine-Water 50%, Water-Lysine 65%, Tryptophan-Water 54%, Water-Tryptophan 65%, Arginine-Water 69% and Water-Arginine 65%). This trend was not observed for non-essential amino acids (i.e., Ornithine, Aspartic Acid, and Glycine), except for aspartic acid (63%) and glycine (50%) when trained as CS-.

**Figure 3.**
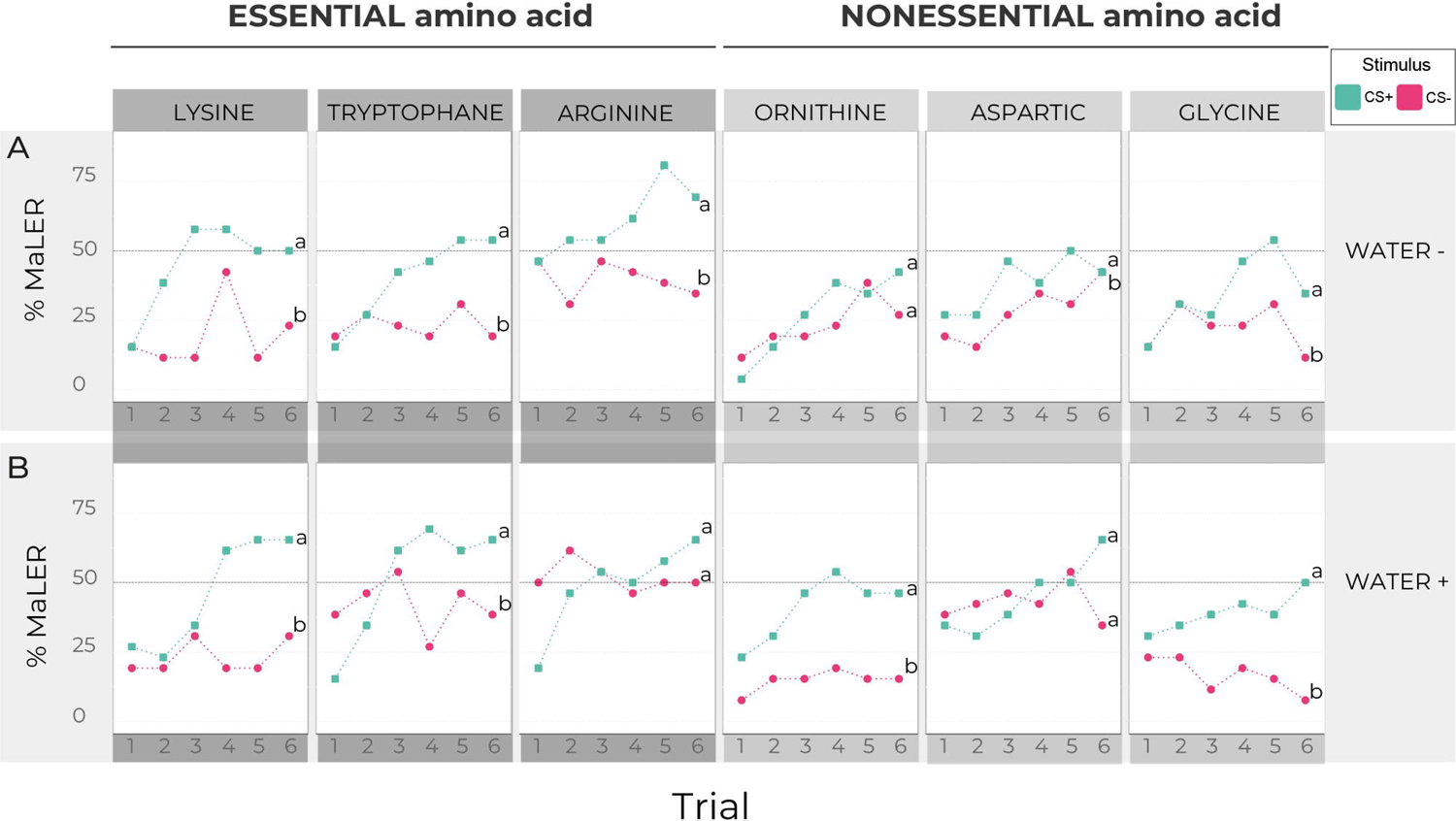
MaLER percentage across trials in Conditioning A: Amino acid vs water. Percentage of MaLER shown by foragers of V*espula germanica* during the 12 trials (six for CS+ and six for CS−) of the 12 sessions in the differential *Conditioning phase* (n=26 for each conditioning session). A) The rewarded stimulus (CS+, green square shape) was an amino acid (i.e., Lysine, Tryptophan, Arginine, Ornithine, Aspartic Acid, or Glycine), and the unrewarded stimulus (CS-, pink circle shape) was the water while in B) is the opposite. Different letters indicate significant differences between the learning curves of each stimulus (p<0.05).

#### Conditioning B: Amino acid vs amino acid

When we analysed the learning curves between rewarded and non-rewarded amino acids (Fig. 4), we observed that wasps could discriminate between amino acids. Notably, they demonstrated the ability to learn and differentiate between two pairs of amino acids, essential and non-essential types. For Lysine and Aspartic Acid learning performance did not depend on the type of rewarded substance (Lysine +/Aspartic acid -: p<0.01, Aspartic acid +/Lysine -: p=0.001). However, in the case of Tryptophan and Glycine, wasps only differentiated between Tryptophan used as CS- and Glycine used as CS+, but not vice versa (Tryptophan +/Glycine -: p=0.098, Glycine +/Tryptophan -: p=0.033).

**Figure 4.**
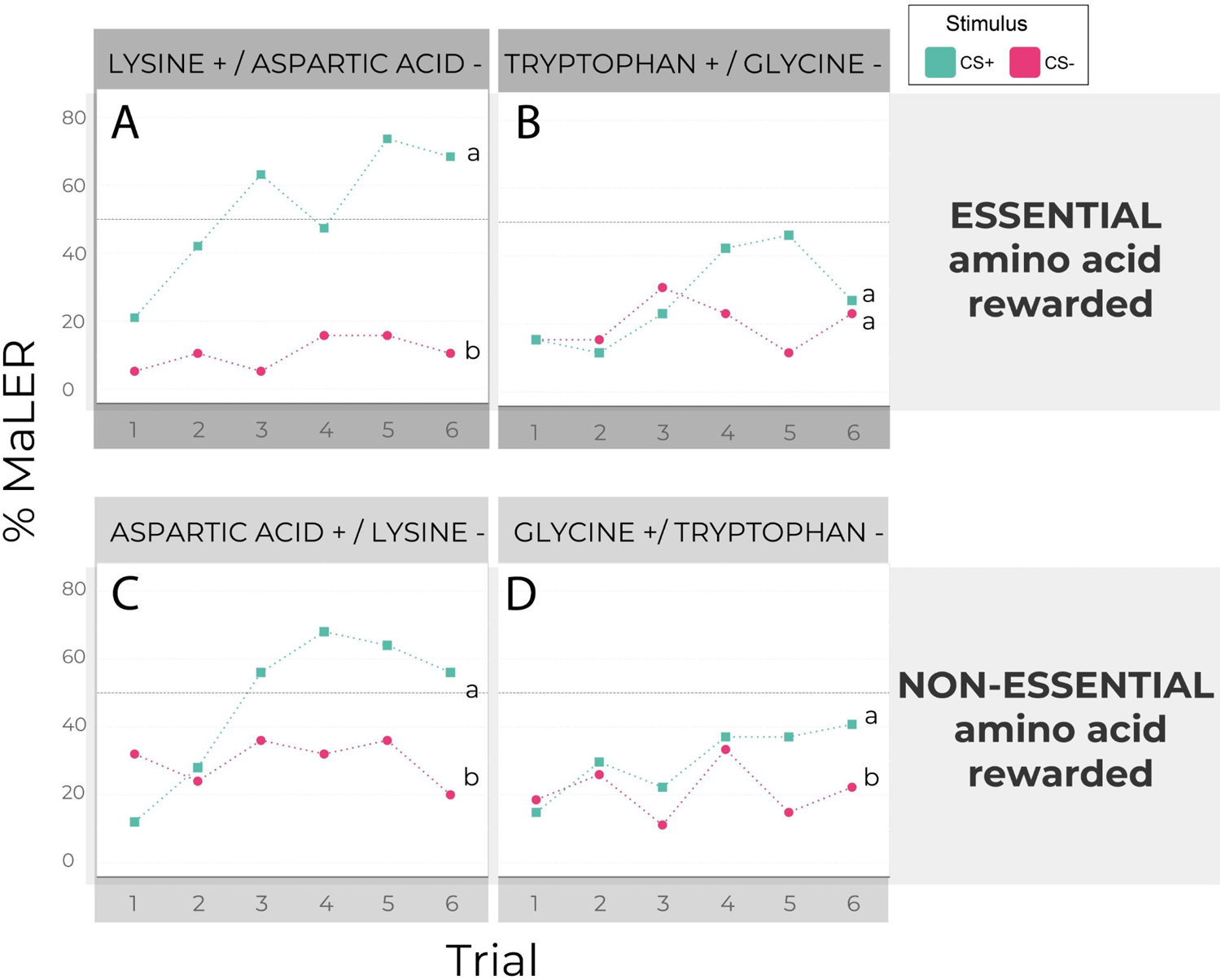
MaLER percentage across trials in Conditioning B: Amino acid vs amino acid. Percentage of MaLER shown by foragers of *Vespula germanica* during the 12 trials (six for CS+ and six for CS−) of the four sessions in the differential *Conditioning phase* (n_Lysine+/Aspartic acid-_=19, n_Aspartic acid+/ Lysine_-=25, n_Tryptophane+/Glycine_-=26, n_Glycine+/ Tryptophane_-=27). The rewarded stimulus (CS+) is represented in a green square shape while the unrewarded stimulus (CS-) is represented in a pink circle shape. A) Lysine (CS+) versus Aspartic acid (CS-), B) Tryptophan (CS+) versus Glycine (CS-), C) Aspartic acid (CS+) versus Lysine (CS-), D) Glycine (CS+) versus Tryptophan (CS-). Different letters indicate significant differences between the learning curves of each stimulus (p<0.05).

### Test phase

#### Conditioning A: Amino acid vs water

The results of the *Test phase* exhibit a high degree of consistency. The memories established for most amino acids remained intact even 30 minutes after the final conditioning event (p_Lys-Wat_=0.006, p_Wat-Lys_=0.041, p_Trp-Wat_=0.04, p_Arg-Wat_=0.01, p_Wat-Orn_=0.01, p_Asp-Wat_=0.014; Fig. 5, green and pink bars). Notably, wasps conditioning with Glycine (as CS+ and CS-) and tryptophan as CS-were unable to discriminate these stimuli from water after the 30-minute interval (p_Gly-Wat_=0.306, p_Wat-Gly_=0.385, p_Wat-Trp_=0.131). Regarding amino acids that failed to be learned during conditioning (Arginine and Aspartic acid, both as CS-), they too were indistinguishable from water after the same 30-minute interval (p_Wat-Arg_=0.857, p_Wat-Asp_=0.0518). However, unexpectedly wasps that initially could not discriminate positively rewarded ornithine from water were able to discriminate both stimuli after 30 minutes (p_Orn-Wat_=0.023).

**Figure 5.**
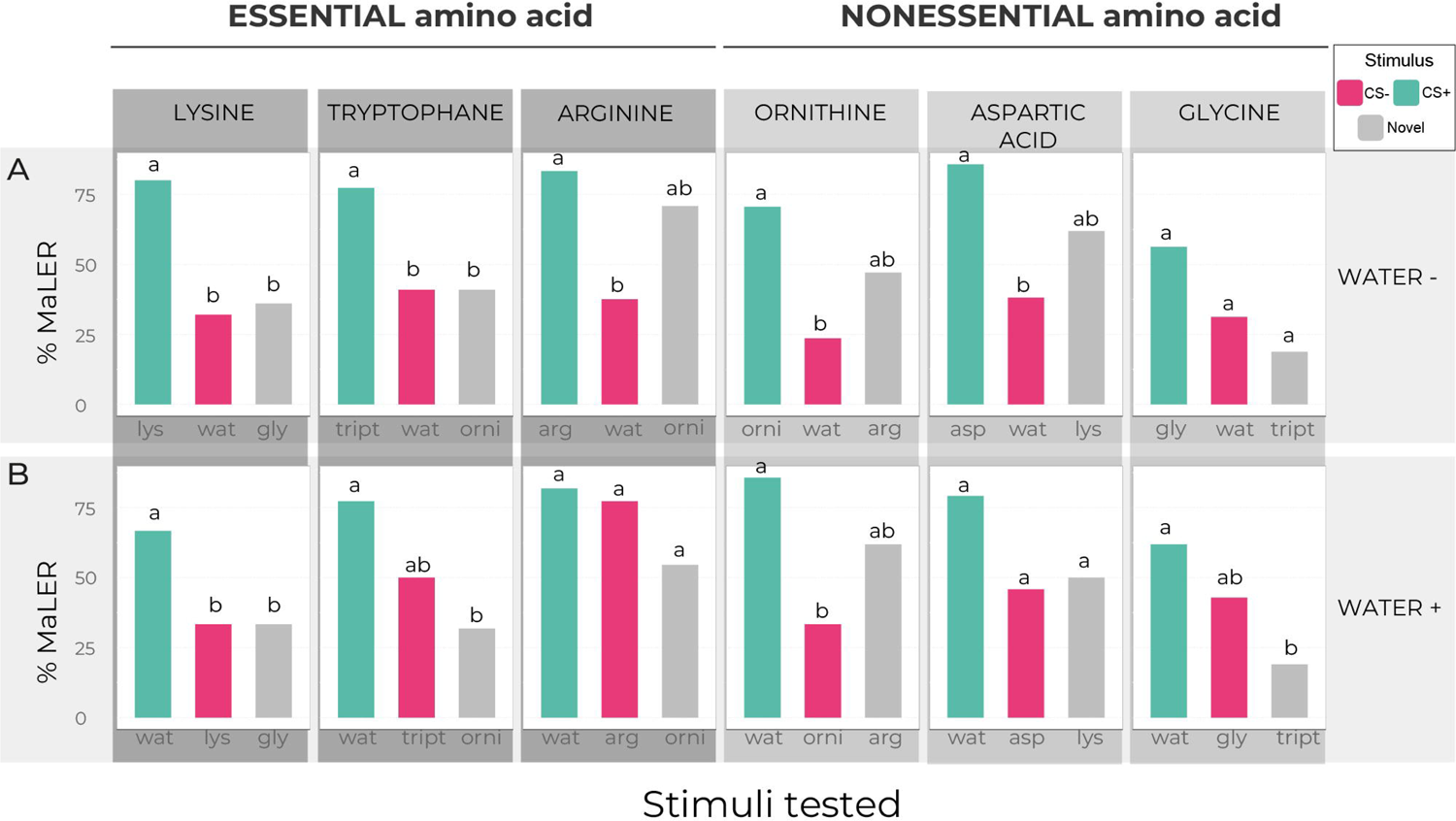
MaLER percentage to learned and novel stimuli during the Test phase for the 12 sessions of Conditioning A. Percentage of MaLER shown by foragers of *Vespula germanica* to the learned stimuli or the novel stimulus (gray bars), during the *Test phase* (n_lys-wat_=25, n_wat-lys_-=24, n_trp-wat_-=22, n_wat-trp_-= 22, n_arg-wat_=24, n_wat-arg_-=22, n_orn-wat_-=17, n_wat-orn_-=21 n_asp-wat_=21, n_wat-asp_-=24, n_gly-wat_-=16, n_wat-gly_-=21). A) In the Conditioning phase, the rewarded stimulus (CS+, green bars) was an amino acid (i.e., Lysine, Tryptophan, Arginine, Ornithine, Aspartic acid, and Glycine) and the unrewarded stimulus (CS-, pink bars) was the water while in B) was the opposite. Different letters indicate significant differences between stimuli (p<0.05).

When testing wasps to a novel stimulus (i.e., an amino acid different from the conditioned one; Fig. 5, green and grey bars), we observed specific patterns of discrimination and generalization. Notably, when wasps were trained with essential amino acids, the acquired learning did not generalize to other non-essential amino acids (p_Lys-Wat_=0.01, p_Wat-Lys_=0.041, p_Trp-Wat_=0.04, p_Wat-Trp_=0.015), except in the case of Arginine, which exhibited generalization when tested with Ornithine (p_Arg-Wat_=0.501, p_Wat-Arg_=0.08). Conversely, when wasps were trained with non-essential amino acids, the acquired learning generalized to other essential amino acids, except for Glycine as CS-(p_Wat-Orn_=0.352, p_Orn-Wat_=0.166, p_Asp-Wat_=0.133, p_Wat-Asp_=0.089, p_Gly-Wat_=0.092, p_Wat-Gly_=0.024).

#### Conditioning B: Amino acid vs amino acid

Wasps conditioned with two amino acids did not show a significant difference in response to the unrewarded amino acid did compared to the response observed with water (negative control) in any of the four conditioning sessions (Lysine +/Aspartic acid - p_Asp-Wat_= 1, Aspartic acid +/Lysine - p_Lys-Wat_= 0.387, Tryptophane +/Glycine - p_Gly-Wat=_ 0.432, Glycine +/ Tryptophane - p_Trp-Wat_= 0.078). Besides, the results for CS+ and CS-responses varied according to the specific pair of rewarded and unrewarded amino acids (conditioning session). When wasps were conditioned to differentiate Lysine as CS+ from Aspartic Acid as CS-(Fig. 6A) or Glycine as CS+ from Tryptophan as CS-(Fig. 6D), the association observed in the conditioning did not persist after 30 minutes; wasps were unable to discriminate between stimuli (p_Lys-Asp_=0.082, p_Gly-Trp_=0.606). When conditioned with Lysine as the CS+, wasps exhibited a tendency for an increased response to this stimulus (Lysine 73% vs. Aspartic acid 27%; differences not significant). Wasps that were conditioned with Tryptophan as the rewarded stimulus and Glycine as the unrewarded one (Fig. 6B), or with Aspartic Acid as the rewarded stimulus and Lysine as the unrewarded one (Fig. 6C), displayed a significantly higher response to the positively rewarded amino acid (Tryptophan or Aspartic Acid) compared to the unrewarded one. Unexpectedly, wasps that during the conditioning event could not discriminate positively rewarded Tryptophan from Glycine were able to discriminate both stimuli after 30 minutes (p_Trp-Gly_=0.006). The memories established remained intact even after a 30-minute post-conditioning period, solely when wasps were conditioned to differentiate Aspartic acid as the CS+ and Lysine as the CS-(p_Asp-Lys_= 0.003).

**Figure 6.**
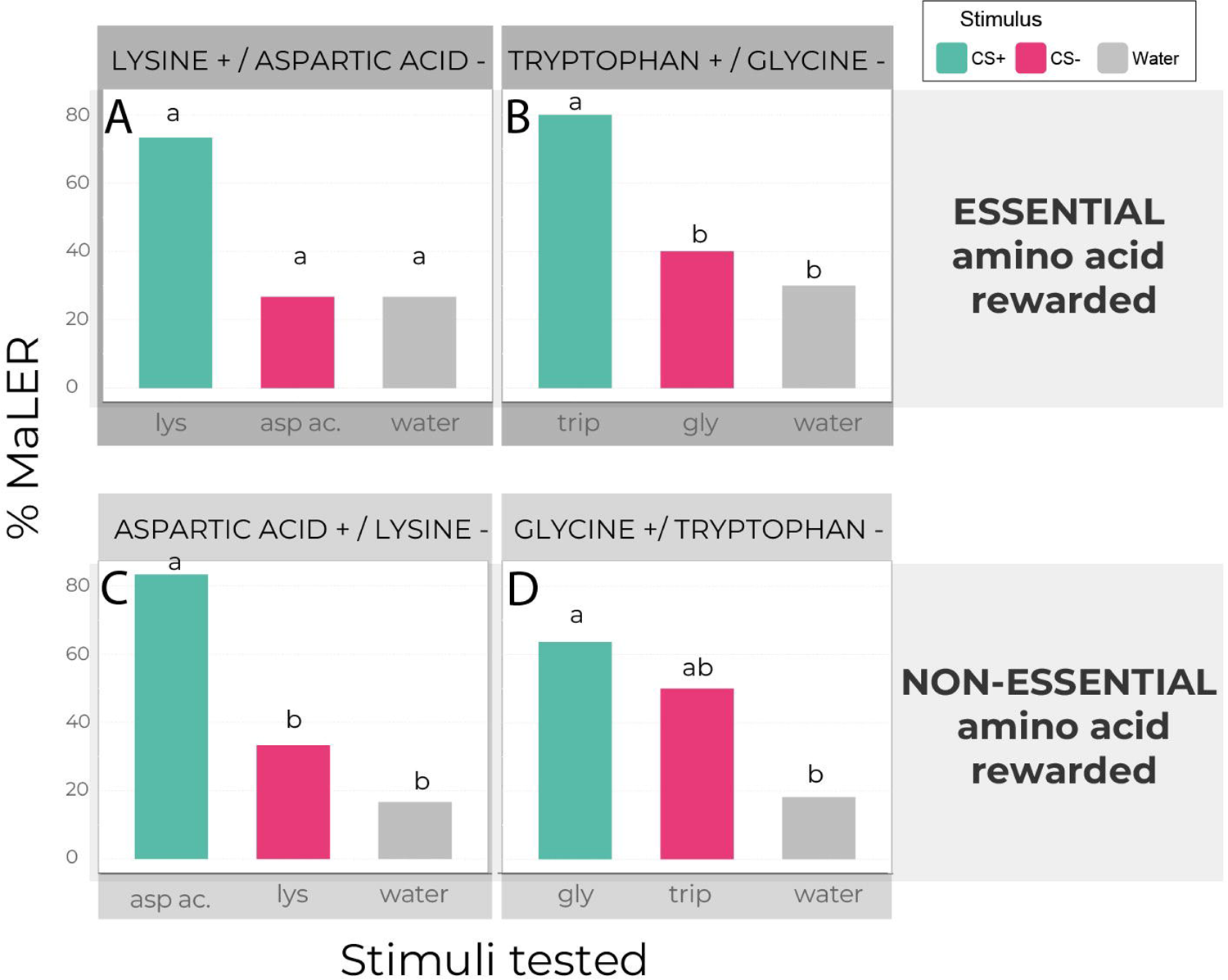
MaLER percentage to learned stimuli and water during the Test phase for the four sessions of Conditioning B. Percentage of MaLER shown by foragers of *Vespula germanica* to the learned stimuli or water (negative control, gray bars), during the *Test phase* (n_Lysine+/Aspartic acid-_=15, n_Aspartic acid+/ Lysine_-=24, n_Tryptophane+/Glycine_-=20, n_Glycine+/Tryptophane_-=22). A or B) In the *Conditioning phase*, the rewarded stimulus (CS+, green bars) was an essential amino acid (i.e., Lysine or Tryptophan) and the unrewarded stimulus (CS-, pink bars) was a non-essential amino acid (i.e., Aspartic acid or Glycine) while in C or D) was the opposite. Different letters indicate significant differences between stimuli (p<0.05).

## Discussion

We studied the chemo-tactile perception and associative learning of *V. germanica* wasps to amino acids. By means of differential conditioning, we show that wasps are capable of perceiving, learning, and discriminating various amino acids from water. In a few cases, discrimination was asymmetrical (i.e., depending on the rewarded stimulus). Indeed, the discriminatory responses between amino acids and water exhibited dissimilarities across the studied stimuli. Discrimination of essential amino acids (Lysine, Tryptophan, and Arginine) was more noticeable than for non-essential amino acids (Ornithine, Aspartic Acid, and Glycine). Also, wasps were able to discriminate between a pair of amino acids, but this behaviour could be asymmetrical depending on which amino acid was rewarded, for example in the session between Tryptophane and Glycine. By means of the *Test phase*, we demonstrate that learned discrimination abilities persisted even after a 30-minute interval, indicating the retention of acquired memories (except for Glycine). Additionally, in certain instances, wasps were able to generalize the acquired information toward novel amino acids. This occurs more frequently when associative learning is with non-essential amino acids and the test is with an essential one. Diverse response abilities exist for amino acids, indicating variations in perception and learning beyond molecular and physiological amino acid attributes. Notably, unlike other Hymenoptera, *V. germanica* wasps can differentiate the qualitative composition of an amino acid solution.

Our findings confirm that *V. germanica* workers, within a cognitive framework, can discriminate some biologically significant amino acids from water. Furthermore, under specific conditions, yellowjacket wasps demonstrated the ability to generalize the learned stimulus to novel amino acids. Notably, when wasps were conditioning with Arginine, they perceived it as similar to Ornithine and vice versa. This pattern may be attributed to the shared involvement of both amino acids in a common metabolic pathway, where Arginine serves as the precursor to Ornithine (Pratavieira et al., 2019). Moreover, Arginine, Ornithine and Lysine - which trigger clear differential learning curves - are amino acids involved in the metabolic transformations of biogenic amines, like putrescine and cadaverine. It has been demonstrated that in worker bees the PER requires the synthesis of substantial quantities of those amines suggesting a key role in the execution of foraging-related behaviours (Pratavieira et al., 2019). Also, these byproducts of decomposing flesh are an attractive signal for certain animals, including scavenging rats and fish, and insects that deposit their eggs on carcasses (Hamana and Matsuzaki, 1984; Heale et al., 1996; Rolen et al., 2003). Thus, based on these facts, we argued that *V. germanica* perceives, learns, discriminates, and generalizes these amino acids because of its scavenging habit. Consequently, it is unsurprising that this species exhibits stronger sensory and perceptual abilities for cues elicited by amino acids compared to non-scavenging hymenopterans (such as bees).

Even though most insects share the need for the same essential amino acids, their perception and relative value could vary between species. Previous studies performed on *A. mellifera* suggest that foragers are unable to perceive and respond to some solutions rich in amino acids (Arenas and Farina, 2012; Carlesso et al., 2021). In contrast, workers of *Bombus terrestris* were able to evaluate the general content of a protein solution, although did not to perceive its composition qualitatively (Ruedenauer et al., 2019). Some authors consider that wasps are the most carnivorous social insects, and as such derive nearly all their protein needs from animal sources (e.g., arthropod and vertebrate prey, and carrion; see Jeanne and Taylor, 2009). Thus, we argue that the discrimination ability of *V. germanica* may be related to their scavenging feeding habits, nutritional requirements, and diet composition of the adult worker wasps. It has recently been demonstrated that, unlike other social insects that collect protein mainly for feeding their brood, adult wasps could metabolize proteins. It has been observed that *V. orientalis* and *V. vulgaris* can digest amino acids in the absence of larvae and use them as metabolic fuel (Teulier et al., 2016; Bodner et al., 2022). Additionally, intestinal proteases were detected in *V. germanica,* indicating the possibility that this species may also be capable of metabolically exploiting these resources (Grogan and Hunt, 1977). To fully understand the role of protein resource consumption in adult individuals it is fundamental to carry further al studies evaluating, for example, their effect on survival and flight capability.

Remarkably, certain amino acids like Arginine and Aspartic acid, exhibited differentiation from water under positive conditioning but not under negative conditioning. When presenting both pure water and water in a solution with an amino acid as stimuli, there is a potential for a confounding effect to emerge, stemming from the shared presence of the same molecule in both stimuli (CS+ and CS-) and thereby disrupting the CS-US contingency, complicating the learning process. Notably, this confounding effect is not observed across all evaluated amino acids. We consider the possibility that the learning experience is intricately linked to the relative salience of both stimuli—water and amino acids. Within this context, simultaneous presentation to the animal may induce perceptual interferences, such as overshadowing or blocking suggesting that, in such instances, only a partial perception of water and amino acids occurs within the compound. Also, stimuli in nature may not appear as isolated, distinct elements. Usually, they are compounds constituted of multiple elements (Mackintosh, 1971). Exploring these cognitive aspects in future studies could yield insightful results.

In conclusion, our study represents a pioneering investigation into the individual perceptual and learning capabilities of *V. germanica* wasps exposed to proteinaceous stimuli in controlled laboratory conditions. We observed varied discriminatory responses among different amino acids and water, with essential amino acids displaying heightened discriminability. The persistence of learned discrimination abilities, and the generalization of acquired information to novel amino acids, highlight the complexity of wasps’ cognitive processes. These findings not only contribute valuable insights into the cognitive dimensions of *V. germanica* but also hold potential implications for pest control strategies. Understanding the learning mechanisms of these wasps could pave the way for the improvement of the control strategies already used (such as in the use of toxic baits) and to develop of innovative approaches in utilizing their cognitive responses for targeted pest management. We also suggest that it would be highly valuable to conduct studies using an integrated approach, merging behavioural ecology with neuroscience, to unravel the mechanisms through which chemo tactile stimuli are processed in the neural systems of social insects. Especially when evaluating other species than the conventional study models, such as pollinating insects, generating a broader perspective. Further research in this direction may offer sustainable and effective solutions for environmentally friendly pest control strategies.

## Acknowledgements

We are grateful to Lic. Romanella Marcellino (IFAB-INTA CONICET) and Dra. Paula Crego (Universidad Nacional del Comahue-CRUB) for providing the amino acids needed for conducting the experiments. The comments of the anonymous referees helped clarify the manuscript.

## Abbreviations

CS: Conditioned stimulus
CS+: Rewarded conditioned stimulus
CS−: Unrewarded conditioned stimulus
+: Rewarded
−: Unrewarded
GLMM: Generalized linear mixed effect model
ITI: Inter trial interval
MaLER: Maxilla labium extension response
PER: Proboscis extension response
US: Unconditioned stimulus
Lys: lysine
Trp: tryptophan
Arg: arginine
Orn: ornithine
Asp: aspartic acid
Gly: glycine

## References

Arenas A & Farina WM (2012). Learned olfactory cues affect pollen-foraging preferences in honeybees, *Apis mellifera*. Animal Behaviour 83(4), 1023–1033. DOI: 10.1016/j.anbehav.2012.01.026

Bell WJ (1990). Searching behaviour patterns in insects. Annual Review of Entomology 35, 447–467. DOI: 10.1146/annurev.en.35.010190.002311

Bitterman ME, Menzel R, Fietz A. & Schafer S (1983). Classical conditioning of proboscis extension in honeybees (*Apis mellifera*). Journal of Comparative Psychology. 97(2), 107–119. DOI: 10.1037/0735-7036.97.2.107

Bodner L, Bouchebti S, Watted O, Seltzer R, Drabkin A & Levin E (2022). Nutrient Utilization during Male Maturation and Protein Digestion in the Oriental Hornet. Biology, 11(2):241. DOI:10.3390/biology11020241

Carlesso, D., Smargiassi, S., Pasquini, E., Bertelli, G., & Baracchi, D. (2021). Nectar non-protein amino acids (NPAAs) do not change nectar palatability but enhance learning and memory in honeybees. Scientific reports, 11(1), 11721. DOI: 10.1038/s41598-021-90895-z

Chapman, R. F. (1998). The insects: structure and function. Cambridge university press.

D’Adamo P & Lozada M (2008). Foraging behaviour in *Vespula germanica* wasp re-locating a food source. New Zealand Journal of Zoology 35 (1), 9–17-DOI: 10.1080/03014220809510099

Dukas R (2008). Evolutionary Biology of Insect Learning. Annual Review of Entomology 53,145–160. DOI: 10.1146/annurev.ento.53.103106.093343

Getz WM & Smith KB (1987). Olfactory sensitivity and discrimination of mixtures in the honeybee *Apis mellifera*. Journal of Comparative Psychology A 160, 239–245. DOI:10.1007/BF00609729

Ghirlanda G & Enquist M (2003). A century of generalization. Animal Behaviour 66, 15–36. DOI:10.1006/anbe.2003.2174

Giurfa M & Sandoz JC (2012). Invertebrate learning and memory: Fifty years of olfactory conditioning of the proboscis extension response in honeybees. Learning and Memory 19, 54–66. DOI: 10.1101/lm.024711.111

Grogan DE & Hunt JH (1977). Digestive proteases of two species of wasps of the genus *Vespula*. Insect Biochemistry 7(3), 191–196. DOI:10.1016/0020-1790(77)90013-0.

Guerrieri F, Schubert M, Sandoz JC & Giurfa M (2005). Perceptual and neural olfactory similarity in honeybees. PLoS Biology. 3, e60. doi:10.1371/journal.pbio.0030060

Guerrieri FJ & d’Ettorre P (2010). Associative learning in ants: Conditioning of the maxilla-labium extension response in *Camponotus aethiops*. Journal of Insect Physiology 56 (1), 88–92. DOI: 10.1016/j.jinsphys.2009.09.007

Gumbert A (2000). Color choices by bumble bees (*Bombus terrestris*): innate preferences and generalization after learning. Behavioural Ecology and Sociobiology 48, 36–43. DOI:10.1007/s002650000213

Hothorn, T., Bretz, F. and Westfall, P. (2008) Simultaneous inference in general parametric models. Biometrical Journal: Journal of Mathematical Methods in Biosciences, 50, 346–363. DOI: 10.1002/bimj.200810425

Hunt, J. H., Baker, I., & Baker, H. G. (1982). Similarity of amino acids in nectar and larval saliva: the nutritional basis for trophallaxis in social wasps. Evolution, 36(6), 1318–1322. DOI: 10.2307/2408164

Jeanne, R. L., & Taylor, B. J. (2009). Individual and social foraging in social wasps. Food exploitation in social insects: ecological behavioral, and theoretical approaches, 53–79.

Käfer H, Kovac H & Stabentheiner A (2012) Resting metabolism and critical thermal maxima of vespine wasps. Journal of Insect Physiology 58(5), 679–689. DOI: 10.1016/j.jinsphys.2012.01.015.

Klowden, M. J. Physiological Systems in Insects. 3rd edn. (Elsevier Science, 2013).

Laska M, Galizia G, Giurfa M & Menzel R (1999). Olfactory discrimination ability and odours structure–activity relationships in honeybees. Chemical Senses 24, 429–443. DOI:10.1093/chemse/24.4.429

Lopatina NG, Zachepilo TH, Kamyshev NG & Chalisova NI. (2017). The influence of combinations of encoded amino acids on associative learning in the honeybee *Apis mellifera* L. Journal of Evolutionary Biochemistry and Physiology 53, 123–128. DOI: 10.1134/S1234567817020045

Lozada M & D’Adamo P (2011). Past Experience: A Help or a Hindrance to Vespula germanica Foragers? Journal of Insect Behaviour 24, 159–166. DOI: 10.1007/s10905-010-9244-6

Mackintosh, N. J. (1971). An analysis of overshadowing and blocking. Quarterly Journal of Experimental Psychology, 23(1), 118–125. DOI: 10.1080/00335557143000121

Marchi, I. L., Palottini, F., & Farina, W. M. (2021). Combined secondary compounds naturally found in nectars enhance honeybee cognition and survival. Journal of Experimental Biology, 224(6), jeb239616. DOI: 10.1242/jeb.239616

Masciocchi M, Mattiacci A, Villacide JM, Buteler M, Porrino AP & Martínez AS (2023). Sugar responsiveness could determine foraging patterns in yellowjackets. Scientific Reports, 13(1), 20448. DOI: 10.1038/s41598-023-47819-w

Matsumoto Y, Menzel R, Sandoz J-C & Giurfa M (2012). Revisiting olfactory classical conditioning of the proboscis extension response in honey bees: a step toward standardized procedures. Journal of Neuroscience Methods, 211, 159–167. DOI:10.1016/j.jneumeth.2012.08.018

Mattiacci, A., Goñalons, C. M., Masciocchi, M., & Corley, J. C. (2023). Gustatory responsiveness in Vespula germanica workers: exploring the interplay between sensory perception and task specialization. Insect Science. DOI: 10.1111/1744-7917.13258

Mattiacci A, Masciocchi M & Corley JC (2021). Flexible foraging decisions made by workers of the social wasp *Vespula germanica* (Hymenoptera: Vespidae) in response to different resources: influence of ontogenetic shifts and colony feedback. Insect Science 29, 581–594, DOI 10.1111/1744-7917.12942

Page Jr RE, Erber J & Fondrk KM (1998). The effect of genotype on response thresholds to sucrose and foraging behaviour of honeybees (*Apis mellifera* L.). Journal of Comparative Physiology A 182, 489–500 (1998). DOI: 10.1007/s003590050196

Pereira AJ, Masciocchi M, Bruzzone O & Corley JC (2013) Field preferences of the social wasp *Vespula germanica* (Hymenoptera: Vespidae) for protein-rich baits. Journal of Insect Behaviour 26, 730–739. DOI: 10.1007/s10905-013-9388-2

Perry, T.W. Animal life-Cycle Feeding and Nutrition; Cunha, T.J., Ed.; Academic Press, Inc.: London, UK, 1984; ISBN 0125520603.

Pietrantuono AL, Requier F, Fernández-Arhex V, Winter J, Huerta G, Guerrieri F. Honeybees generalize among pollen scents from plants flowering in the same seasonal period. Journal of Experimental Biology 222 (21), jeb201335. DOI:10.1242/jeb.201335

Pratavieira, M., da Silva Menegasso, A. R., Roat, T., Malaspina, O., & Palma, M. S. (2019). In situ metabolomics of the honeybee brain: The metabolism of l-arginine through the polyamine pathway in the proboscis extension response (PER). Journal of proteome research, 19(2), 832–844. DOI: 10.1021/acs.jproteome.9b00653

Raveret Richter M (2000). Social Wasp (Hymenoptera: Vespidae) Foraging Behaviour. Annual Review of Entomology 45:121–150. DOI: 10.1146/annurev.ento.45.1.121

Robazzi Bignelli Valente Aguiar JM, Roselino AC, Sazima M. & Giurfa M (2018). Can honey bees discriminate between floral-fragrance isomers? Journal of Experimental Biology, 221, jeb180844. doi:10.1242/jeb.180844

Ruedenauer FA, Leonhardt SD, Lunau K & Spaethe J (2019). Bumblebees are able to perceive amino acids via chemotactile antennal stimulation. Journal of Comparative Physiology A 205, 321–331. DOI: 10.1007/s00359-019-01321-9

Teulier L, Weber J-M, Crevier J & Darveau C-A (2016). Proline as a fuel for insect flight: enhancing carbohydrate oxidation in hymenopterans. Proceedings of the Royal Society B: Biological Sciences. 283,20160333. DOI:10.1098/rspb.2016.0333

